# Toward comprehensive functional analysis of gene lists weighted by gene essentiality scores

**DOI:** 10.1101/2021.04.26.441450

**Authors:** Rui Fan, Qinghua Cui

## Abstract

Gene functional enrichment analysis represents one of the most popular bioinformatics methods for annotating the pathways and function categories of a given gene list. Current algorithms for enrichment computation such as Fisher’s exact test and hypergeometric test totally depend on the category count numbers of the gene list and one gene set. In this case, whatever the genes are, they were treated equally. However, actually genes show different scores in their essentiality in a gene list and in a gene set. It is thus hypothesized that the essentiality scores could be important and should be considered in gene functional analysis. For this purpose, here we proposed WEAT (https://www.cuilab.cn/weat/), a weighted gene set enrichment algorithm and online tool by weighting genes using essentiality scores. We confirmed the usefulness of WEAT using two case studies, the functional analysis of one aging-related gene list and one gene list involved in Lung Squamous Cell Carcinoma (LUSC). Finally, we believe that the WEAT method and tool could provide more possibilities for further exploring the functions of given gene lists.

## INTRODUCTION

Gene functional enrichment analysis represents one class of the most popular tools in bioinformatics (1,2). Within a large group of methods, the hypergeometric test is one of the most frequently used ones and it has been widely used and continuously improved in many online tools including DAVID Bioinformatics Resources (3).

However, the current algorithms for gene functional analysis (e.g. hypergeometric test) are based on a hypothesis that all the genes in one gene set are identical, which is not the true cases in the various biological processes. In fact, different genes have different contributions to one biological process. For example, essential genes are normally more important than non-essential genes in molecular functions. Therefore, one critical problem is that, for any two gene lists with the same count number of hitting genes within a given gene set, current algorithms will show the same significance. But in fact, the significances of the two input gene lists should be different. That is, the list whose hitting genes with greater essentiality scores should be more important than the other list. However, all of the current algorithms fail to address the above issues.

To address the above issue, here we proposed a weighted functional enrichment method based on hypergeometric test by introducing a prior weight to each gene. We first collected a group of scores representing the gene essentiality. Then we developed Weighted Enrichment Analysis Tool (WEAT, https://www.cuilab.cn/weat/) algorithm and tool to perform weighted functional enrichment analysis. Finally, we showed the usefulness of WEAT using two case studies.

## METHODS AND MATERIALS

### Conventional Algorithms for Gene Functional Enrichment Analysis

Hypergeometric test and binomial test have been widely accepted for gene functional enrichment analysis (4). The above two types tests are quite similar and the major difference is the sampling with or without replacement. Thus, the conventional methods for gene functional enrichment analysis is described as follows. Suppose *A* is the intersection set of the inputted gene list and one known gene set. Let *k* be the size of *A*, that is k = |A|. Normally, hypergeometric test is based on hypergeometric distribution, and it is to estimate the probability of X > *k* intersection genes in a specific gene set, which is defined as follows:

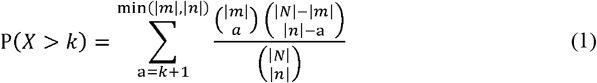

where N is the set of genes in the background, m denotes the inputted gene list, n is a specific known gene set. Hence, we can obtain the significance of each gene set from the probability calculated by equation (1).

### Weighted Algorithm for Gene Functional Enrichment Analysis

Here we presented the algorithm for weighted gene functional enrichment analysis, which is an extension of the hypergeometric test by replacing the gene counts with the sum score of those genes, which is described as follows:

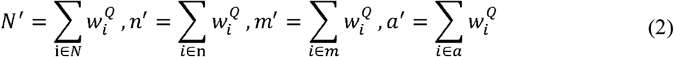

where W_i_ represents the weight for gene i and Q denotes the scaling factor of scores. Nevertheless, introducing the scores into the hypergeometric test violates the discreteness of hypergeometric distribution. Because the calculation of hypergeometric distribution is based on the calculation factorial, which only works for positive integers. Hence, to solve this problem, we introduce the Gamma function as the natural extension of factorial calculation, which is defined as follows:

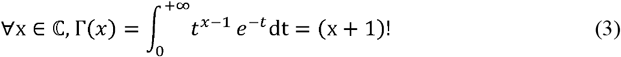

Gamma function can extend the domain of factorial function from positive integers to all complex numbers. Here we only use the positive real part. We then calculate the probability of weighted algorithm with positive real numbers defined as follows:

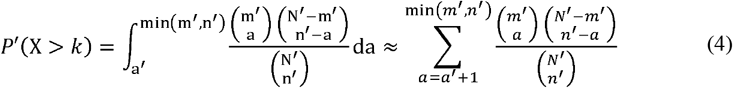

The discrete hypergeometric distribution will become a continuous distribution when applying the Gamma function on it. Therefore, the calculation should change from summation to integration. However, for the simplicity and consistency of the calculation, we still use the discrete form to estimate the probability as defined in equation (4). It should be noted that if all the gene weights equal to one, the weighted algorithm will reduce to the conventional algorithms as shown in equation (1).

Traditional methods (5,6) always take the scores into ranks or round the scores into integers to prevent violation of discreteness, while it scaling up some of *N, m, n*, and *a*, which will also cause deviations from the original values of probability because of the scaling but not weighting. For example, if we scale up all genes to have a score of 2 rather than 1 (count), the expected result should have exactly the same p-value but it is not.

### Score normalization

To keep the consistency between the conventional methods and the weighted method, normalization is needed to keep all the scores have the mean value of one and greater than zero as well, which is defined as follows:

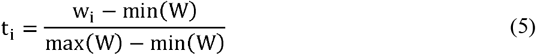

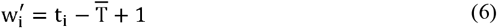

where *W*_*i*_ represents the original weight of the i^th^ gene, W = [w_1_,w_2_, …,w_N_] is the original weight matrix, t_i_, denotes the pre-normalized weight of gene i, T = [t_t_,t_2_, …,t_N_] is the pre-normalized weight matrix, 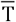 represents the mean value of T, 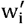 is the normalized weights. As a result, the normalized weights have the mean value of one, which makes the weighted methods comparable to unweighted one.

### Inverse Document Frequency score for distinct categories

Given a large number of gene sets, the frequency of one gene turns up in different sets has a very big difference. Thus, there can be a very intuitive hypothesis: a gene frequently involved in various gene functional terms may cause less effect on individual sets. In natural language processing (NLP), there is a popular concept and algorithm named Inverse Document Frequency (IDF) which is used to evaluate the importance of a word among documents. That is, the more one word shows in other documents, the less important this word will be in any document. Hence, the calculation of the IDF of one gene can be defined as follows:

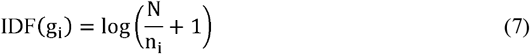

where N is the total number of terms in one category (refer to the concept of corpus in NLP), n_i_ is the count of different terms (refer to the concept of document in NLP) where gene i is included.

### Log proportion normalization

IDF calculation works well for gene sets but not for gene expression profiles among tissues. And, our purpose is trying to extract the tissue-specific genes in a reasonable way. Hence, to evaluate the importance of each gene in different tissues, a log proportion normalization of expression profile is applied, which defined as follows:

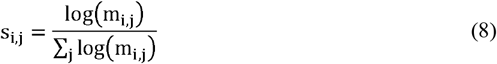

where m_i,j_ represents the gene counts of the i^th^ gene in the j^th^ tissue. Therefore, s, j can be interpreted as the importance score of a specific gene i in different tissues.

### Gene sets and gene essentiality scores

For gene sets, we collected some common gene annotation sets for functional enrichment analysis including Kyoto Encyclopedia of Genes and Genomes (KEGG, https://www.kegg.jp/), Gene Ontology (GO) Resource (7), and gene sets in Enrichr (8). In addition, we collected and processed five groups of gene essentiality scores including evolutionary conservation, expression profiles of specific tissues, gene importance score, protein-protein interaction (PPI) score and IDF of gene set as shown in Table 1.

**Table 1.**
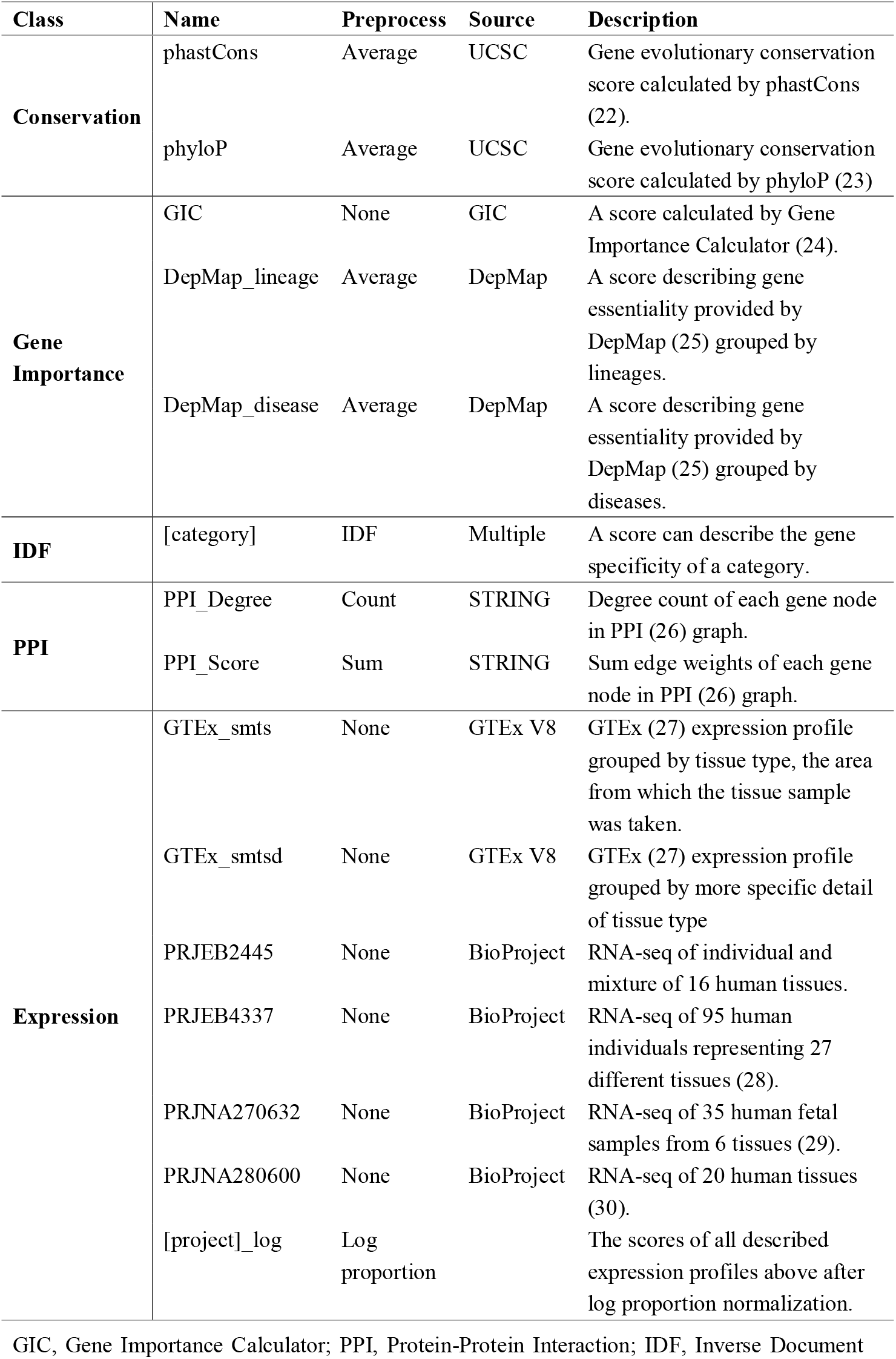
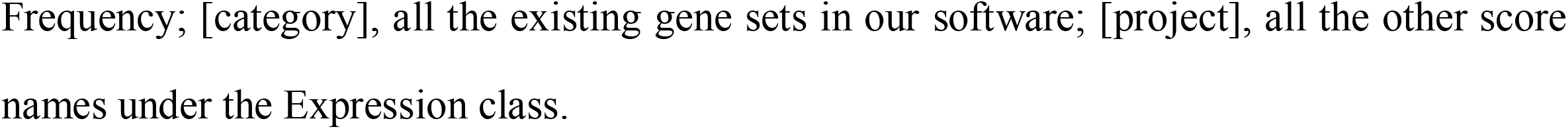
List of gene essentiality scores used in WEAT.

### Web server construction

The web server is built using the Django framework as the back-end for processing and calculation. For the front-end, we are using Bootstrap for web framework design, bootstrap-table for table illustration, Ploty.js for figure visualization, and JQuery for application logic. All the algorithms are implemented in Python using packages Scipy, Numpy, and Pandas. The web server is available at https://www.cuilab.cn/weat/.

## RESULTS

### Overall design and Web server

The basic workflow of weighted gene functional enrichment analysis (WEAT) is shown in Figure 1. In the knowledge part, WEAT collected and curated 200 gene functional categories (140 for Homo Sapiens), 471,296 gene sets (315,285 for Homo Sapiens) and 654 types of gene essentiality scores (563 for Homo Sapiens). Then, the user needs to input a gene list, for example, the deregulated genes from a study of omics. Next, WEAT will calculate the p-value for both the conventional gene functional enrichment analysis methods and the weighted gene functional enrichment analysis method. After that, a comparison will be performed between the results of the two methods. Meanwhile, the gene sets which are only significant in the weighted methods will be highlighted.

**Figure 1.**
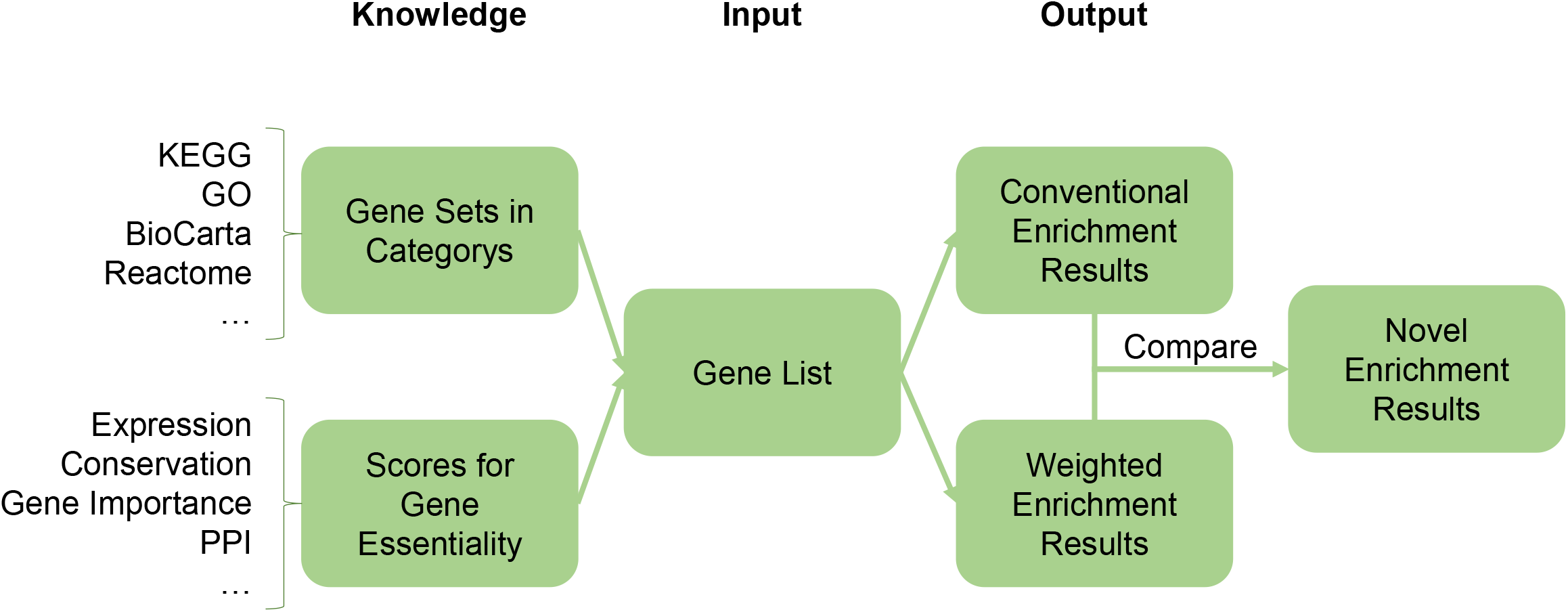
The framework of the weighted gene functional enrichment analysis.

As shown in Figure S1-2, the web interface of WEAT is designed following the above workflow. Firstly, users should select the species and then input a gene list. And then users need to choose a type of gene essentiality score. Meanwhile, WEAT also provides a choice for users to upload their own gene essentiality scores. (Figure S1). Next, after choosing the scale factor Q, users can click the “Run” button to submit the task. On the brief result page (Figure S2), users can perform analyses for any category of gene sets. Meanwhile, users can search, sort, export, and visualize their results.

### Case Studies

#### The Prion pathway is involved in the aging process in human brain

It is known that one common feature of the ageing brain is the age-associated periventricular white matter lesions (PVL), which reflects white matter injury. It is thus important to dissect the functional pathways or terms involved in PVL. For doing so, we applied WEAT to a gene expression dataset of brain PVL caused by aging (GEO: GSE157363) (9). By using the instructions of the dataset, we identified 1715 differentially expressed genes (p-value ≤ 0.05, |log2(fold-change)| ≥ 1.2). Next, we selected the brain gene expression in GTEx as the gene essentiality score. The results on the BioCarta pathway category are shown in Figure 2 (Table S1 for detail information).

**Figure 2.**
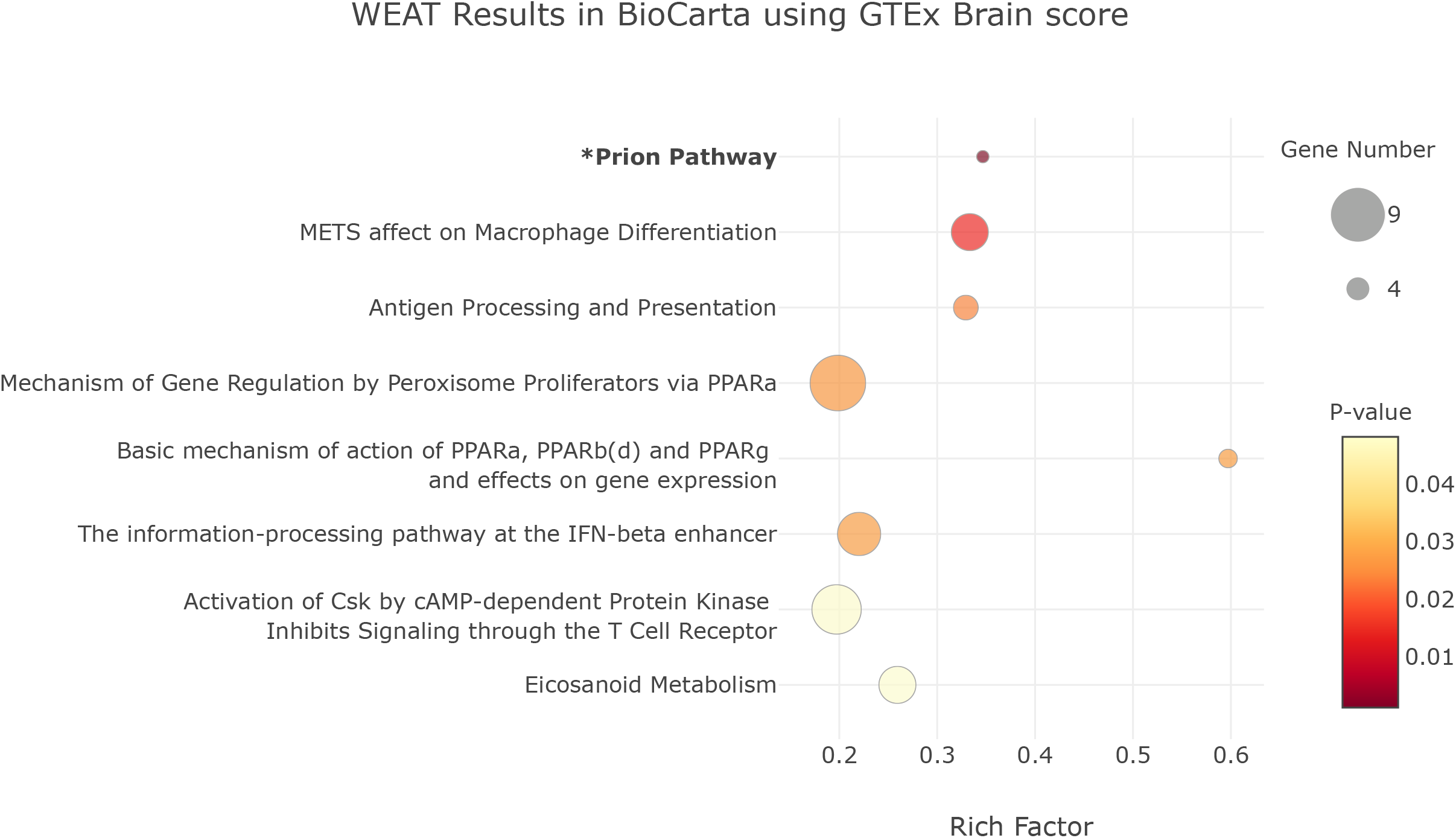
WEAT analysis results of the deregulated genes in age-associated periventricular white matter lesions (PVL) in the BioCarta signaling pathway category using GTEx brain essentiality score. Rich Factor = (sum score of hit genes) / (sum score of all genes in this set)

It should be noted that the Prion pathway shows the greatest statistical significance in the weighted method while is not significant in the conventional ones. There are only two hit genes (GFAP and DNAJB6) between the input gene list and the Prion pathway gene set (Figure 3). Therefore, the conventional methods failed to identify the Prion pathway as significance, whereas the GFAP gene showed a very high normalized gene essentiality score (7.81) in brain tissue, resulting in the greatest significance of this pathway. The conventional methods cannot identify this term because they treated GFAP and other genes equally. In fact, GFAP, the glial fibrillary acidic protein, is highly related to brain and neural function. It was reported that the cellular prion protein (PrP) is greatly associated with Alzheimer’s and Parkinson’s diseases, which are indeed highly related with brain aging and lesions (10,11). This finding supported that WEAT can uncover some important patterns missed by the conventional methods.

**Figure 3.**
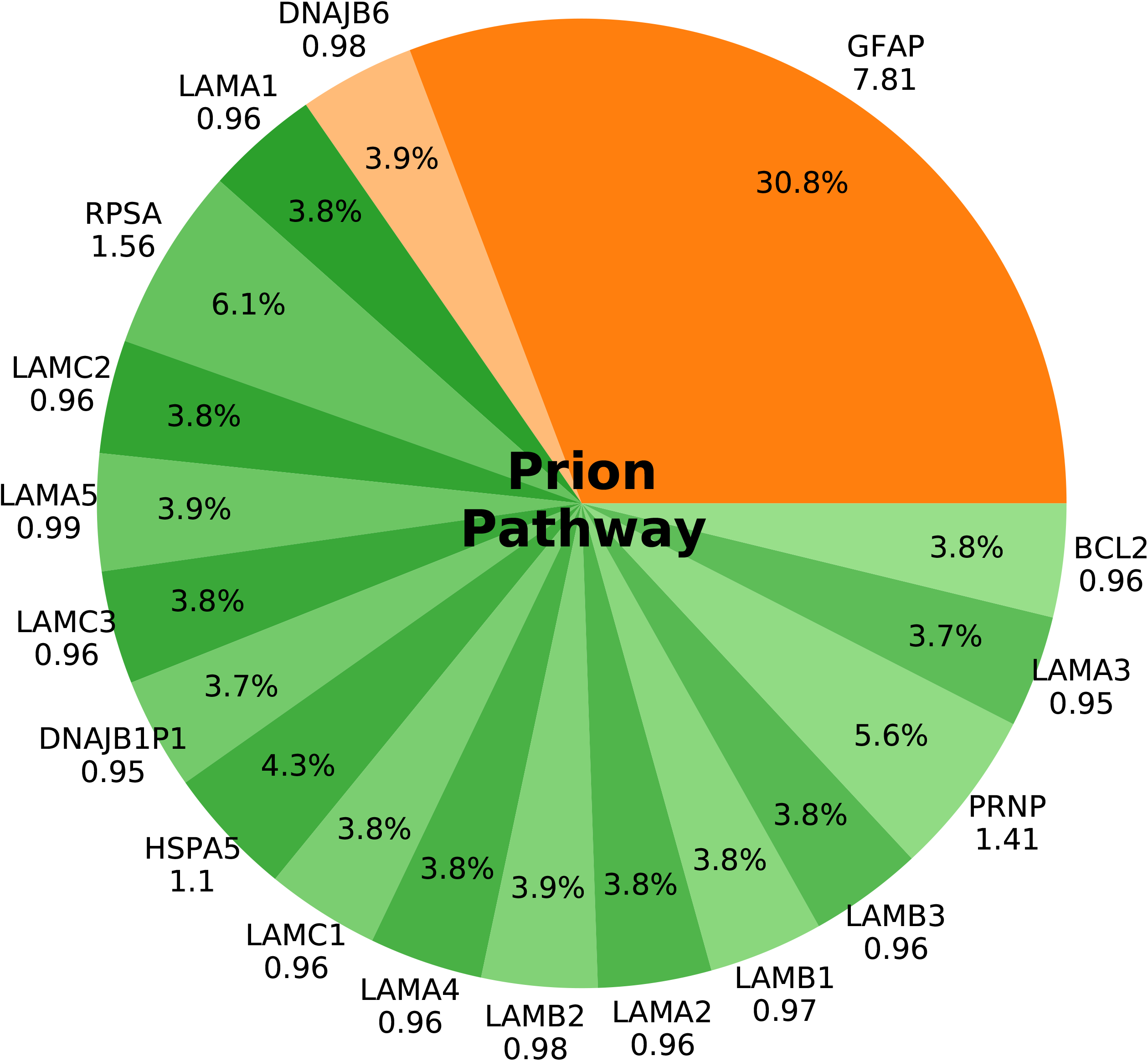
The normalized gene essentiality scores of genes in the Prion pathway by the WEAT analysis of deregulated genes in age-associated periventricular white matter lesions (PVL). The hit genes are shown in orange and the missed genes are shown in green.

### Lung Squamous Cell Carcinoma

Lung Squamous Cell Carcinoma (LUSC) is the most common histologic subtype of non-small cell lung cancer (NSCLC), highly associated with cigarette smoking (12), with poor clinical prognosis (13). Exploring the functional pathways involved in LUSC is important for understanding its mechanisms and finding new therapeutic targets. Here we downloaded the LUSC gene expression data from The Cancer Genome Atlas (TCGA) (14) including 502 tumors and 49 tumor adjacent tissues. Using the edgeR (15) package, we identified 773 differentially expressed genes. For comparison, we used different gene essentiality scores and focused on the KEGG pathways. The novel terms identified by WEAT are shown in Figure 4 (Table S2 for detail information).

**Figure 4.**
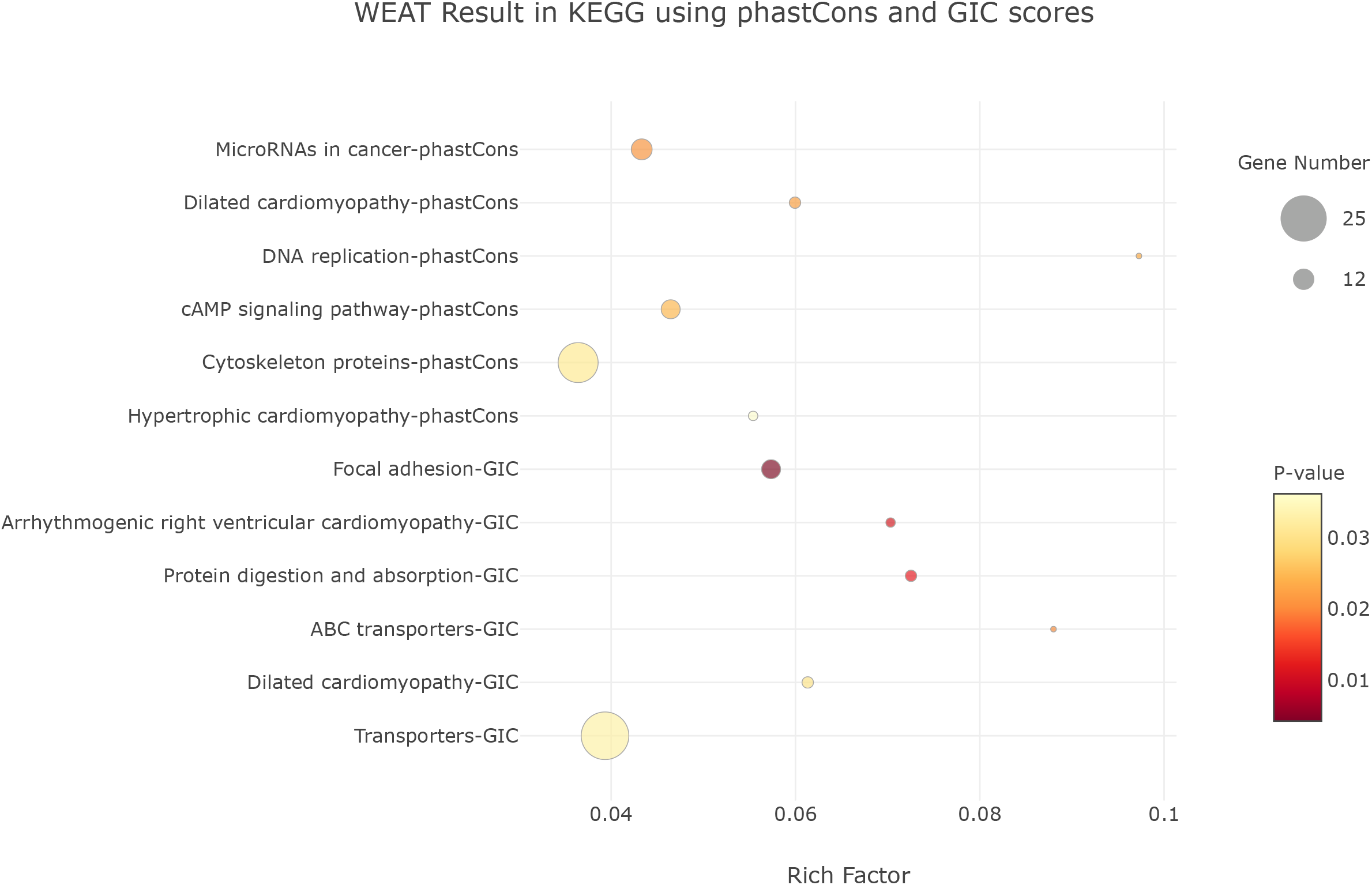
The KEGG pathways identified only by WEAT analysis results but not by the conventional methods for the deregulated genes in Lung Squamous Cell Carcinoma (LUSC) using different gene essentiality scores (phastCons\ score \and GIC score). Rich Factor = (sum score of hit genes) / (sum score of all genes in this set)

As a result, we found that different gene essentiality scores have different preferences for terms without loss of plausibility. For example, there was a patient reported as LUSC with severe cardiac dysfunction similar to dilated cardiomyopathy (16) which was successfully identified by WEAT using the phastCons score. Similarly, a case with lung cancer was found to be related to the dilated phase of hypertrophic cardiomyopathy (17) which was identified by WEAT using the gene importance calculator (GIC) score. Other terms including the disorder of DNA replication (18) and cytoskeleton proteins (19) using the phastCons scores, focal adhesion (20) and ABC transporters (21) using the GIC scores, are indeed the basal pathways of cancer that were not identified by the conventional methods.

### Different scores prefer different terms

As discussed above, different gene essentiality scores have different tendencies on identifying functional terms. Thus, to further investigate the preference of the gene essentiality scores, we calculated the variance of gene essentiality scores for each functional term in the GO biological process (GOBP) as an example. Given that if the gene essentiality scores in a gene set could be greatly different from each other, this set may play a more important role under the situation of the selected score, here we only focus on the terms with greatest variance (Figure S3-5).

As a result, different essentiality scores indeed produce preferences to specific functional terms. As shown in Figure S3, for essentiality scores derived from GTEx gene expression, all the tissue expressed based scores prefer to essential functional terms such as hydrogen peroxide biosynthetic process and electron or proton transport (right part of Figure S3). Moreover, each tissue also prefers to tissue-specific functional terms (left part of Figure S3). For example, the essential score derived from the blood prefer to gas-related functions such as nitric oxide transport and oxygen transport. Meanwhile, WEAT also provided normalized GTEx essential scores. The normalization of the GTEx essential scores will eliminate the common functional terms and focus on the tissue-specific terms (Figure S4). For essentiality scores of PPI degree and PPI scores, they are very similar and have a higher preference to the functional terms that the hub genes are involved in. The conservation scores including phastCons and phyloP show preference to the evolutionary conservative pathways such as embryonic development functions; The GIC score prefers to functional terms related to essential biological processes; The IDF score are more likely to identify functional terms with unique genes. In all, different essentiality scores can make the enrichment result subtle and hence the weighted method is able to identify different reasonable functional terms by using different scores.

## DISCUSSION

Gene functional enrichment analysis is one of the most popular ways for annotating the functions of a given gene list. However, the genes in one input gene list and one gene set are equally treated for both hypergeometric test and binomial test, which are the basic algorithms for the conventional methods. This would lead to biased results because genes are actually different in essentiality.

In this study, we introduced a weighted gene functional enrichment analysis method, WEAT, by considering the above truth that different genes have different essentiality in one biological process. We further implemented the WEAT algorithm, developed the online tool, and confirmed its usefulness by two case studies. As a result, WEAT shows its capacity in exploring important gene functional terms which the conventional methods failed to dissect. WEAT could be extended to various applications. For example, if we choose a gene expression essentiality score different from the tissue of the input case, we may find some functional terms involved in the crosstalk between the two tissues. Hence, this provides a potential way to find inter-organ interactions. Moreover, the scale factor Q is positive in the current WEAT algorithm, which means higher gene essentiality leads to greater weight in a gene functional term. While it can be negative, which could be useful to explore the trivial functional terms when using IDF scores. In addition, WEAT could be improved in a number of aspects, for example, more gene sets and gene essentiality scores in the knowledge base, and more species will be annotated in the future. Finally, the profile-level two-phenotype gene set enrichment analysis could be improved using the idea of WEAT.

## DECLARATIONS

### Ethics approval and consent to participate

Not applicable.

### Consent for publication

Not applicable.

### Availability of data and materials

The datasets generated and analyzed during the current study are available on our website at https://www.cuilab.cn/weat/. The datasets of case studies are available at https://www.ncbi.nlm.nih.gov/geo/query/acc.cgi?acc=GSE157363 and https://portal.gdc.cancer.gov/projects/TCGA-LUSC.

### Authors’ contributions

Q.C. conceived the project. R.F. collected the data, designed the algorithm, performed the analyses and built the web application. R.F. and Q.C. wrote and edited the manuscript and approved the final manuscript.

---

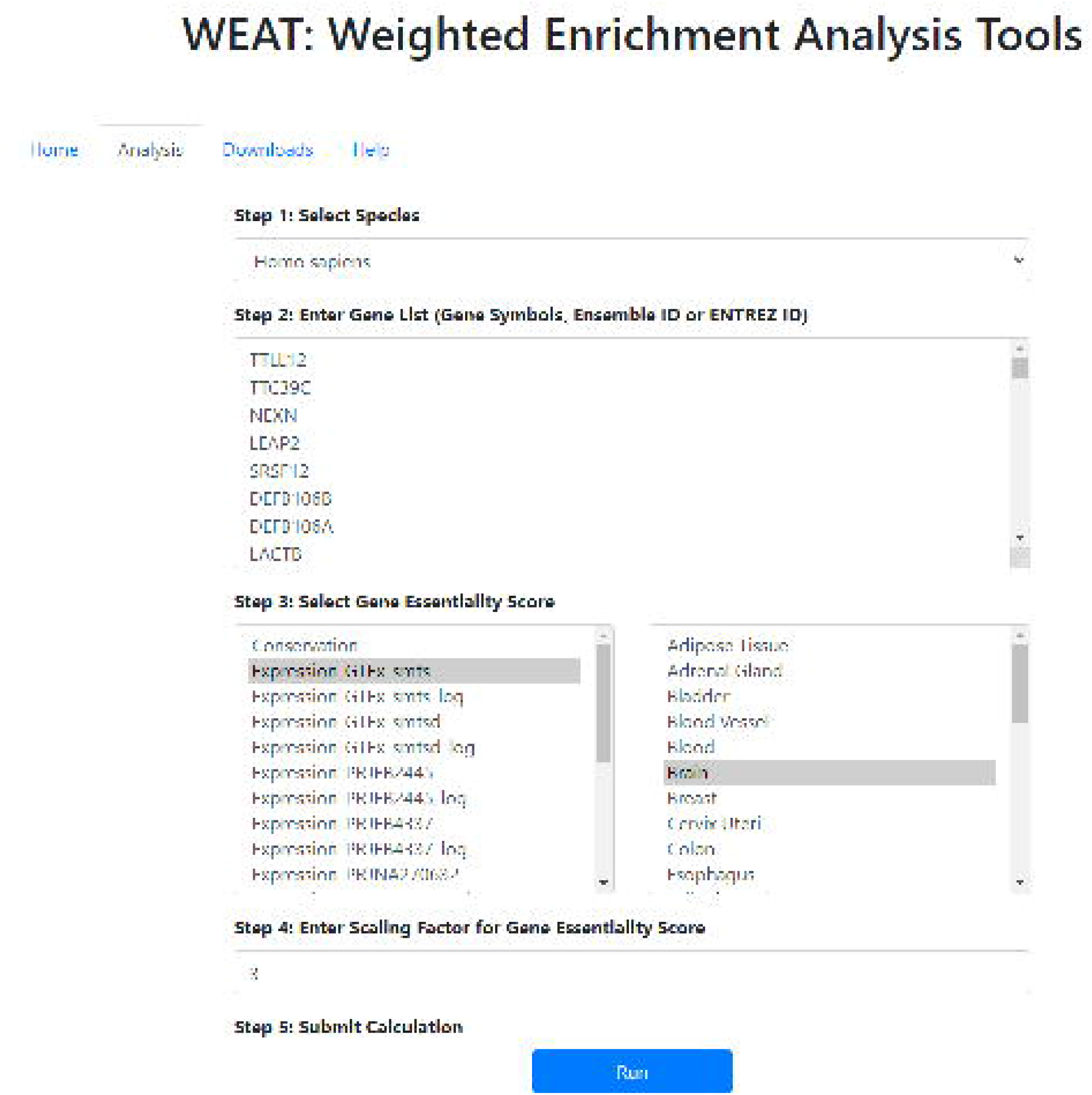

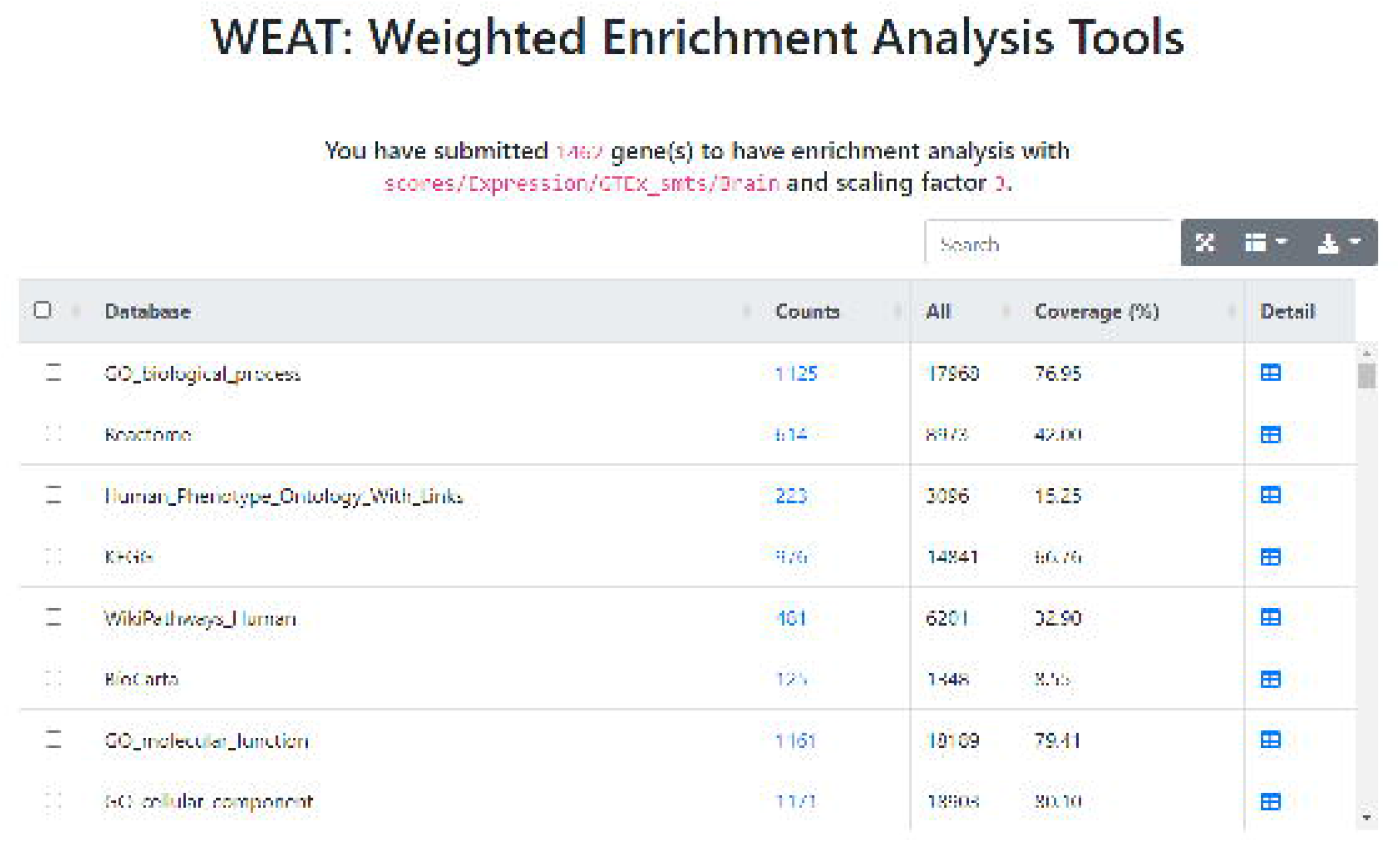

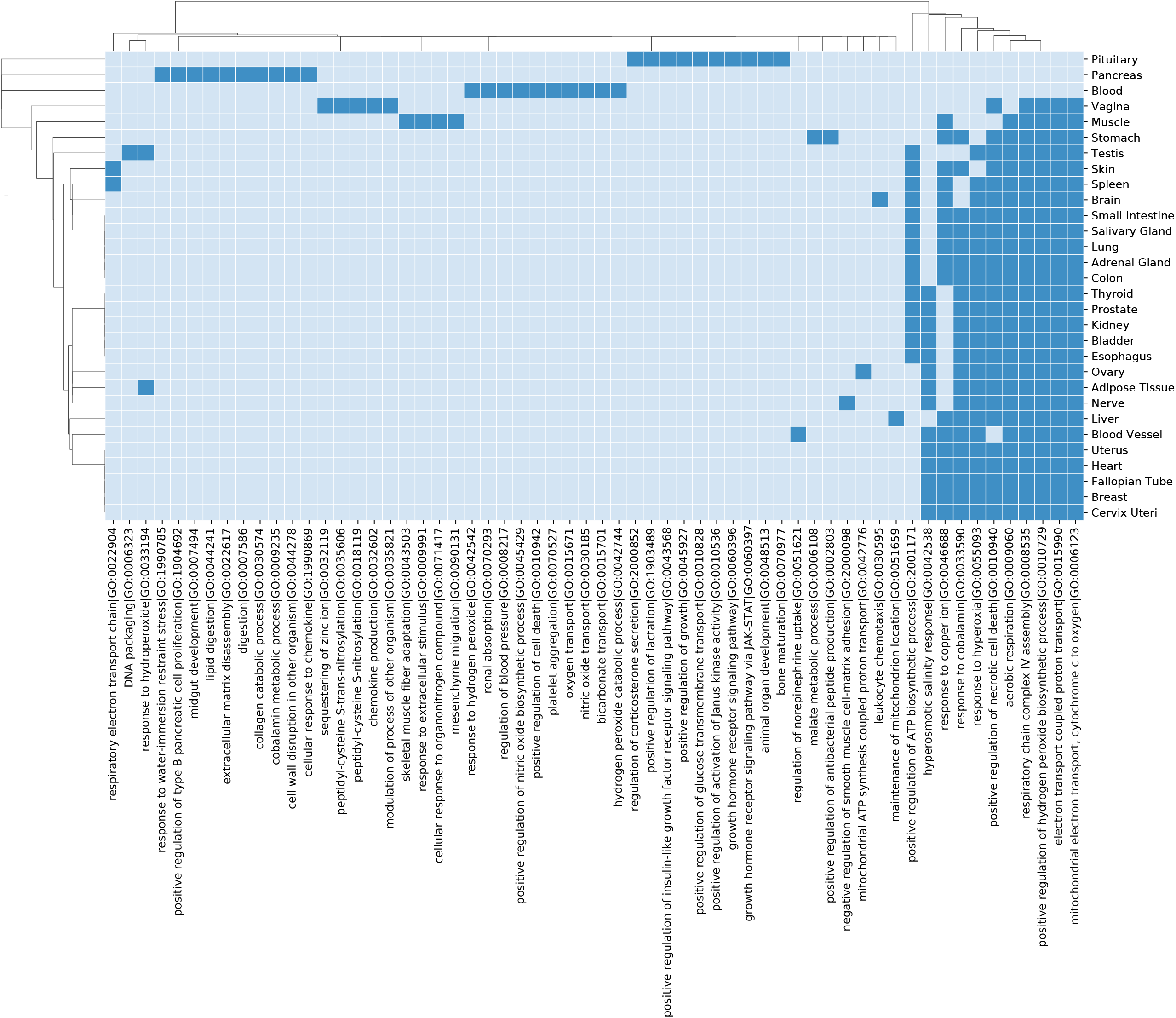

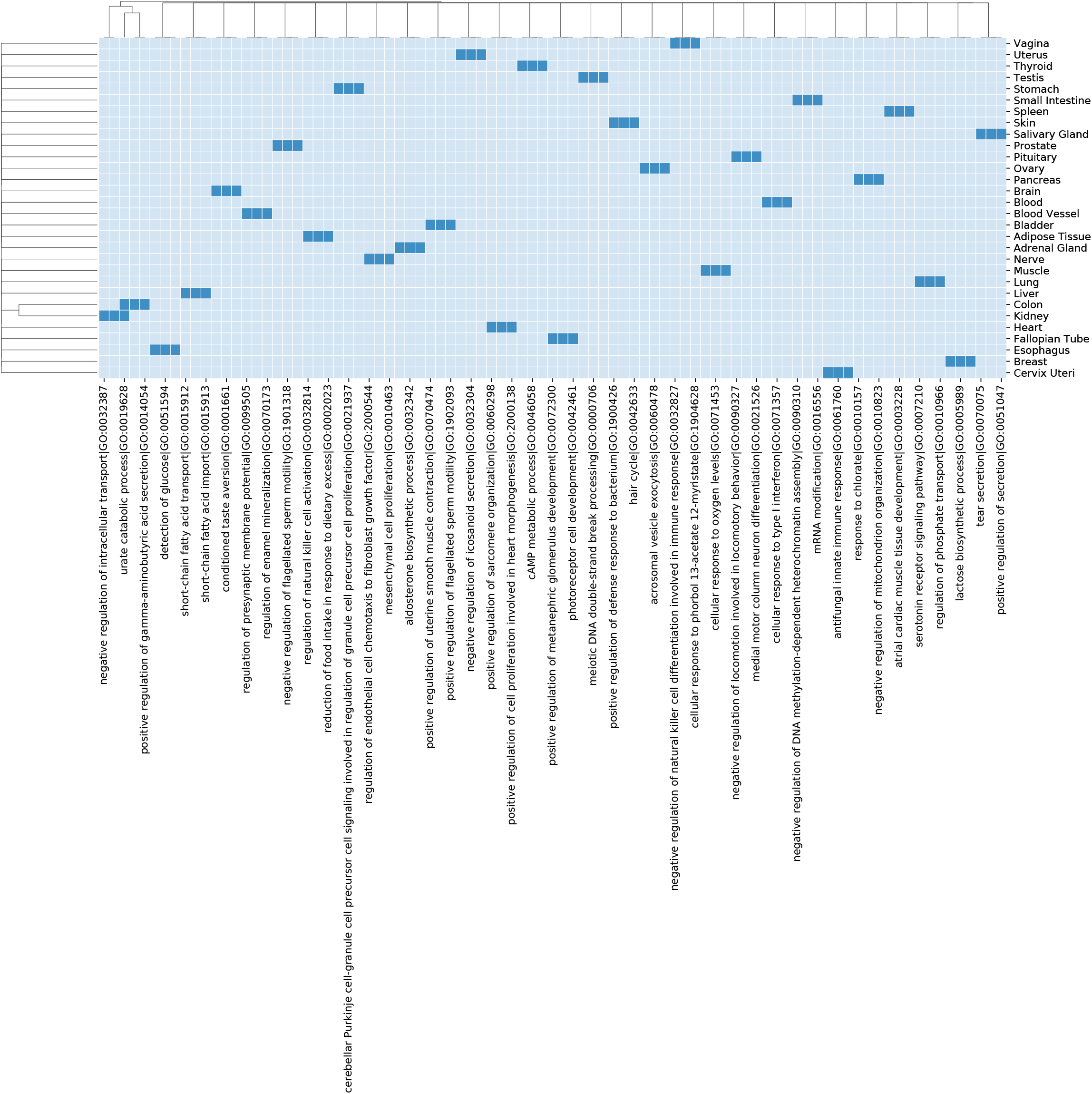

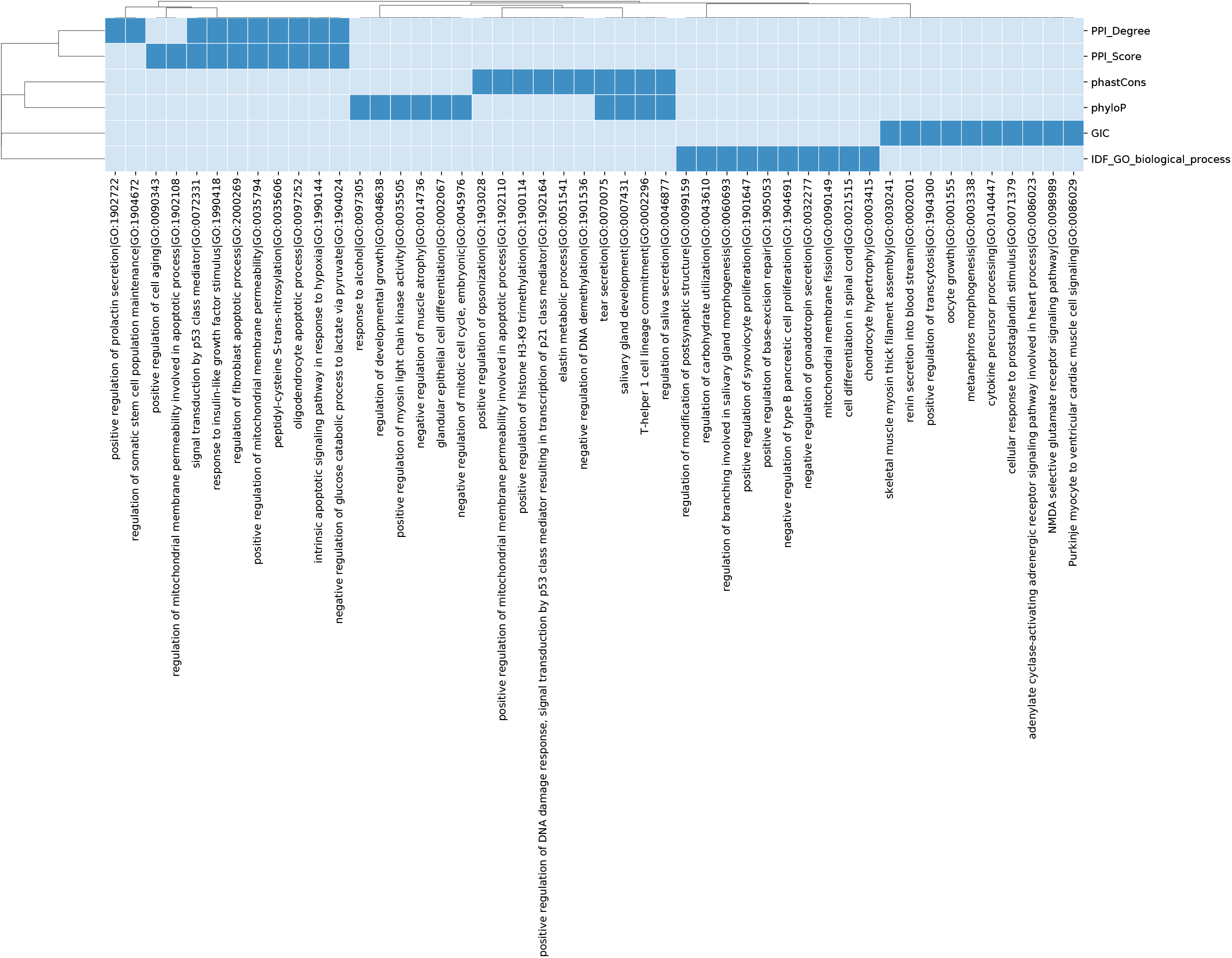

