## Supplementary material for "Toward comprehensive functional analysis of gene lists weighted by gene essentiality scores": Table S1 and Table S2

**Table S1. WEAT analysis results of the deregulated genes in age-associated periventricular white matter lesions (PVL) in the BioCarta signaling pathway category using GTEx brain essentiality score.**

| **Term** | **Counts** | **OR** | **Scores** | **WOR** | **P-value** | **WP-value** |
| --- | --- | --- | --- | --- | --- | --- |
| ***Prion Pathway** | 2/18 | 1.23 | 8.79/25.36 | 4.79 | 0.5090 | 0.00139 |
| METS affect on Macrophage Differentiation | 6/18 | 5.09 | 5.96/17.88 | 4.3 | 0.0041 | 0.0152 |
| Antigen Processing and Presentation | 4/12 | 5.02 | 4.31/13.09 | 4.67 | 0.0194 | 0.0241 |
| Mechanism of Gene Regulation by Peroxisome Proliferators via PPARa | 9/52 | 2.13 | 10.53/53.04 | 2.27 | 0.0448 | 0.0267 |
| Basic mechanism of action of PPARa, PPARb(d) and PPARg and effects on gene expression | 3/5 | 15.01 | 2.89/4.84 | 18.5 | 0.0068 | 0.0272 |
| The information-processing pathway at the IFN-beta enhancer | 7/29 | 3.24 | 7.01/31.83 | 2.75 | 0.0137 | 0.0273 |
| Activation of Csk by cAMP-dependent Protein Kinase Inhibits Signaling through the T Cell Receptor | 8/43 | 2.32 | 8.53/43.25 | 2.22 | 0.0393 | 0.0481 |
| Eicosanoid Metabolism | 6/23 | 3.58 | 5.75/22.17 | 2.92 | 0.0152 | 0.0484 |

OR, Odds Ratio; WOR, Weighted Odds Ratio; WP-value, Weighted P-value; *, novel term.

**Table S2. The KEGG pathways identified only by WEAT analysis results but not by the conventional methods for the deregulated genes in Lung Squamous Cell Carcinoma (LUSC) using different gene essentiality scores (phastCons score and GIC score).**

| Score | Term | Counts | OR | Scores | WOR | WP-value |
| --- | --- | --- | --- | --- | --- | --- |
| phastCons | MicroRNAs in cancer | 11/310 | 1.3 | 16.17/373.34 | 1.82 | 0.0214 |
|  | Dilated cardiomyopathy | 6/96 | 2.36 | 8.51/141.96 | 2.43 | 0.0225 |
|  | DNA replication | 3/36 | 3.21 | 4.45/45.75 | 3.93 | 0.0237 |
|  | cAMP signaling pathway | 10/216 | 1.72 | 13.40/288.46 | 1.92 | 0.0251 |
|  | Cytoskeleton proteins | 21/534 | 1.46 | 27.58/757.40 | 1.51 | 0.0313 |
|  | Hypertrophic cardiomyopathy | 5/90 | 2.08 | 7.09/127.97 | 2.35 | 0.0363 |
| GIC | Focal adhesion | 10/201 | 1.86 | 16.75/292.06 | 2.2 | 0.00444 |
|  | Arrhythmogenic right ventricular cardiomyopathy | 5/77 | 2.46 | 8.05/114.48 | 2.83 | 0.0105 |
|  | Protein digestion and absorption | 6/103 | 2.19 | 8.67/119.53 | 2.73 | 0.0126 |
|  | ABC transporters | 3/45 | 2.52 | 5.33/60.57 | 3.4 | 0.0204 |
|  | Dilated cardiomyopathy | 6/96 | 2.36 | 8.61/140.39 | 2.29 | 0.0303 |
|  | Transporters | 25/676 | 1.37 | 25.32/643.76 | 1.53 | 0.0329 |

WP-value, Weighted P-value; GIC, Gene Importance Calculator.

**Figures**

**Figure S1. The “Analysis page” of the WEAT tool.**

**Figure S2. The “Brief result page” of the WEAT tool.**

**Figure S3. The preference of different GTEx gene essentiality scores in gene ontology biological process (GOBP) category.** The top 10 GOBP terms with greatest variance of gene essentiality scores for each specific tissue are colored as deep blue.

**Figure S4. The preference of different normalized GTEx gene essentiality scores in gene ontology biological process (GOBP) category.** For the convenience of visualization, here only the top 3 GOBP terms with greatest variance of gene essentiality scores for each specific tissue are colored as deep blue.

**Figure S5. The preference of different gene essentiality scores (PPI degree, PPI score, phastCons score, phyloP score, GIC score, and IDF of GOBP score) in gene ontology biological process (GOBP) category.** The top 10 GOBP terms with greatest variance of gene essentiality scores for each specific tissue are colored as deep blue.
